# Striatal compartmentalization and clustering of different subtypes of astrocytes is altered in the zQ175 Huntington’s disease mouse model

**DOI:** 10.1101/2021.11.29.470488

**Authors:** Taylor G Brown, Mackenzie Thayer, Nicole Zarate, Rocio Gomez-Pastor

## Abstract

Huntington’s disease (HD) is a devastating neurodegenerative disease that primarily affects the striatum, a brain region that controls movement and some forms of cognition. Dysfunction and loss of medium spiny neurons of the striatum is accompanied by astrogliosis (increased astrocyte density and pathology). For decades, astrocytes were considered a homogeneous cell type, but recent transcriptomic analyses revealed astrocytes are a heterogeneous population classified into multiple subtypes depending on the expression of different gene markers. Here, we studied whether three different striatal astrocyte subtypes expressing glutamine synthetase (GS), glial fibrillary acidic protein (GFAP), or S100 calcium-binding protein B (S100B) are differentially altered in HD. We conducted a comparative immunofluorescence analysis in the striatum of WT and the heterozygous zQ175 HD mouse model and found that the expression and abundance of GFAP+ and S100B+ astrocytes increased in zQ175 mice, while GS+ astrocytes showed no alteration. We then explored whether there was a differential spatial distribution of any of these subtypes within the striatum. We developed a systematic brain compartmentalization approach and found that while GS+ and S100B+ astrocytes were more homogeneously distributed throughout the striatum in zQ175 mice, GFAP+ astrocytes preferentially accumulated in the dorsomedial and dorsolateral striatum, which are regions associated with goal-directed and habitual behaviors. Additionally, GFAP+ astrocytes in zQ175 mice showed increased clustering, a parameter that indicates increased proximity and that is associated with localized inflammation and/or neurodegeneration. Our data suggest a differential susceptibility in both increased density and striatal compartmentalization of different subtypes of astrocytes in zQ175. These results highlight new potential implications for our understanding of astrocyte pathology in HD.

## Introduction

Huntington’s disease (HD) is a devastating neurodegenerative disease that manifests as progressive motor, cognitive, and psychiatric impairments (Roos, 2010). A CAG (glutamine) triplet expansion in the Huntingtin gene (*HTT)* produces a dysfunctional mutant protein (mHTT) that is prone to misfolding and aggregation (The Huntington’s Disease Collaborative Research Group, 1993). mHTT preferentially affects medium spiny neurons (MSNs) in the striatum and leads to degeneration of this brain region. Anatomical analyses showed that within the striatum the dorsal striatum is severely affected (Bates et al., 2015; Morigaki & Goto, 2017), although the reason for such region selectivity is uncertain.

HD is also characterized by astrogliosis, defined as an increased density of astrocytes, and astrocytic dysfunction (Verkhratsky et al., 2019; Vonsattel et al., 1985). Astrocytes are glial cells with a variety of homeostatic, synaptic, and neuroprotective functions (Araque et al., 1999; Eroglu & Barres, 2010; Verkhratsky et al., 2019). Specifically, an increased density of astrocytes and an upregulation of the astrocytic glial fibrillary acidic protein (GFAP), which is considered a canonical marker of astrocyte reactivity (Buffo et al., 2008; Escartin et al., 2021; Hol & Pekny, 2015; Kamphuis et al., 2012), is seen in postmortem brains of patients with HD (Selkoe et al., 1982; Vonsattel et al., 1985). Additionally, astrocyte dysfunction, in terms of ion homeostasis, Ca^2+^ signaling, and neurotransmitter clearance, is a key factor in both the onset and progression of HD symptoms (Al-Dalahmah et al., 2020; Diaz-Castro et al., 2019; Khakh et al., 2017). When mHTT is explicitly expressed in mouse astrocytes there is a progressive disruption of astrocytic glutamate transport and the manifestation of some HD-like phenotypes (Faideau et al., 2010), while reduction of mHTT in astrocytes of BACHD mice partially improved neuronal excitability and motor behavior (Wood et al., 2019). These studies support the fundamental role of astrocytes in HD pathology.

Contrary to previous knowledge, recent transcriptomic studies have shown that astrocytes are a heterogeneous group of cells that differ transcriptionally and morphologically by brain region (Batiuk et al., 2020; Hasel et al., 2021; Khakh & Deneen, 2019; Ohlig et al., 2021; Sosunov et al., 2014) and can have multiple, distinct states of reactivity depending on the environmental perturbations (Al-Dalahmah et al., 2020; Hasel et al., 2021; Liddelow et al., 2017; X. Yu et al., 2020). Single cell RNA-seq (scRNA-seq) evidence in postmortem HD patient brains demonstrated distinct subtypes of astrocytes that are categorized based on the expression of different astrocytic markers and implied multiple response states in HD (Al-Dalahmah et al., 2020). Therefore, our previous understanding of the contribution of astrocytes to HD may be incomplete. These new studies raised new questions as to whether different subtypes of astrocytes may be preferentially affected in HD and whether they play a differential role in HD pathology.

Given the clear evidence for astrocyte heterogeneity and their involvement in HD, we sought to determine whether different subtypes of astrocytes present different states of astrogliosis and whether they shared a common or distinct spatial distribution throughout the striatum. We studied three subtypes of astrocytes, each expressing a different astrocyte marker: glutamine synthetase (GS), S100 calcium binding protein B (S100B), and GFAP in the brain of wildtype (WT) and symptomatic zQ175 (HD) mice. We conducted immunofluorescence (IF) analyses and 2D imaging to assess density, spatial distribution, and clustering of different astrocytes subtypes. Our results demonstrate a selective increase in astrocyte density in zQ175 mice in S100B+ and GFAP+ astrocytes while no changes in GS+ astrocytes were found. In addition, increased density and clustering of GFAP+ astrocytes were restricted to the dorsomedial and dorsolateral regions of the striatum contrary to S100B which presented an overall increase throughout the striatum. Overall, our data demonstrated a differential alteration in the density and spatial distribution of different astrocyte subtypes in the striatum of HD mice that could have a direct impact in understanding their distinct roles in pathology.

## Results

### Astrocyte subtypes exhibit differences in density and distribution throughout the striatum in zQ175 mice

Increased astrogliosis is a hallmark of HD. As increasing evidence for multiple subtypes of astrocytes with potential distinct functions arises (Al-Dalahmah et al., 2020; Hasel et al., 2021; Khakh & Deneen, 2019), the necessity for determining whether specific subtypes of astrocytes are differentially involved in HD becomes more essential. To investigate whether HD-related astrogliosis affects different astrocyte subtypes, we utilized three different immunofluorescence markers of astrocytes, GS, S100B, and GFAP, which each correspond to a different cluster of astrocytes as characterized by recent scRNA-seq studies (Al-Dalahmah et al. 2020). GS is traditionally viewed as a pan-astrocyte marker, S100b is considered a marker of mature astrocytes and expression of GFAP is associated with reactive astrocytes (Anlauf & Derouiche, 2013; Buffo et al., 2008; Escartin et al., 2021; Hol & Pekny, 2015; Kamphuis et al., 2012). We performed immunofluorescence on coronal brain sections from WT and zQ175 mice at 12 months. At this age, zQ175 mice show robust motor deficits, and therefore this is considered a symptomatic time point (Heikkinen et al., 2012; Menalled et al., 2012; Yu and Zarate et al., 2020). We assessed astrocyte density, calculated as the number of cells per unit area, in three specific striatal locations (subregions) that correspond to dorsomedial (dm), ventromedial (vm) and ventrolateral (vl) (**Fig. 1A**) using three antibody markers (**Fig. 1B**).

**Fig. 1.**
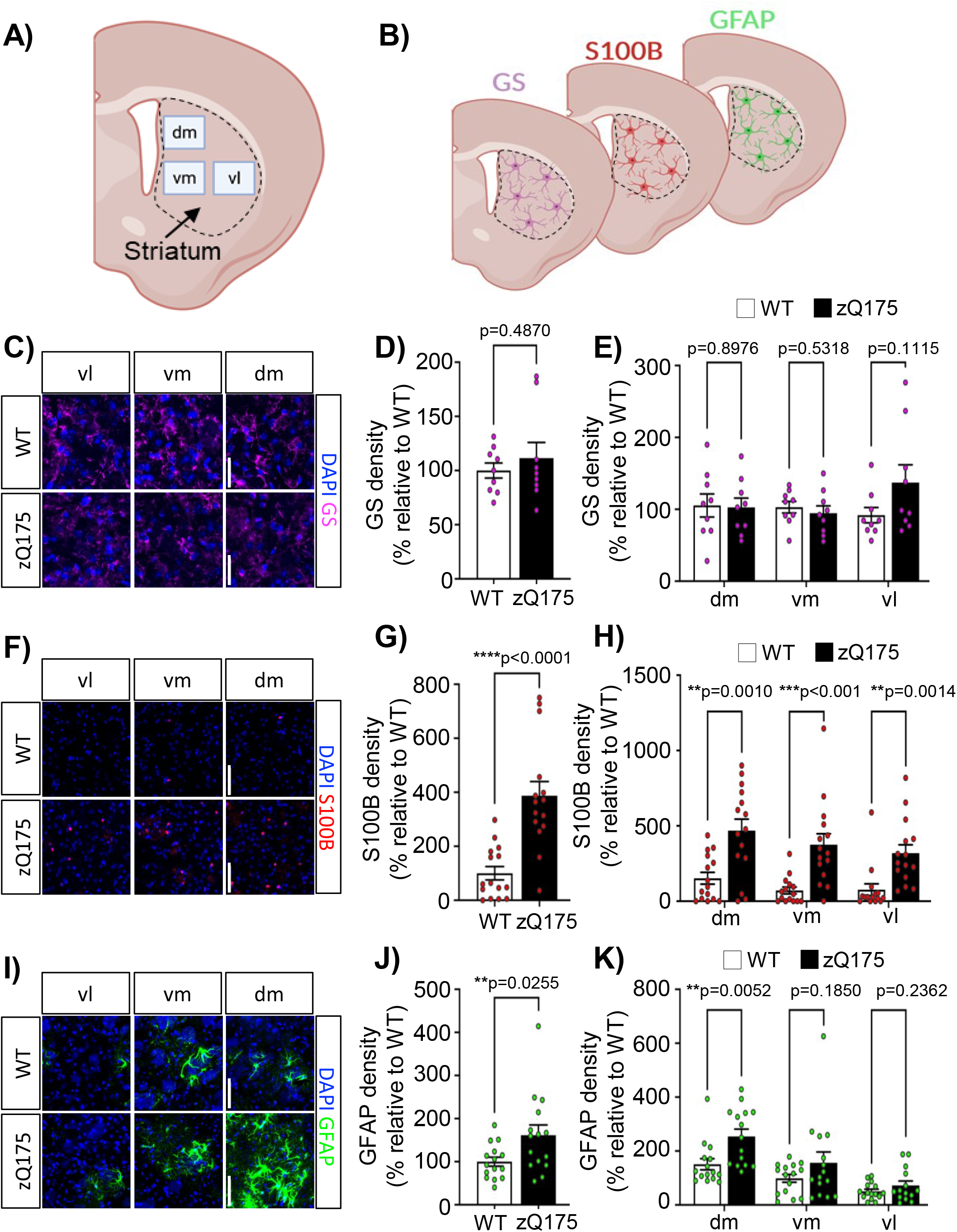
Selective astrogliosis of specific astrocyte subtypes in the striatum in zQ175 mice. (**A**) Representative coronal section of the mouse brain showing the striatum. Rectangles representing dorsomedial (dm), ventromedial (vm), and ventrolateral (vl) regions are marked. Images were taken in each of the three regions and density was calculated for each. Image was created with BioRender.com. (**B**) Diagram of experimental design. Coronal slices were stained for either GS, S100B, or GFAP and striatal regions were imaged. Created with Biorender.com. (**C**) Representative GS (magenta) immunostaining. (**D**) GS density in whole striatum and (**E**) in individual striatal regions was calculated from C and relativized to WT (n=9 slices from n=3 mice/genotype). (**F**) Representative S100B (red) immunostaining. (**G**) S100B density in whole striatum and (**H**) in striatal regions (n=15 slices from n=5 mice/genotype). (**I**) Representative GFAP (green) immunostaining. (**J**) GFAP density in whole striatum and (**K**) in striatal regions (n=15 slices from n=5 mice/genotype). Images were taken at 10X zoom in WT and zQ175 mice at 12 months. DAPI stains nuclei. Scale bar, 50 μm. Density in the whole striatum is calculated as the sum between dm+vl+vm. Error bars denote mean ± SEM. Un-paired Student’s t test. *p<0.05, **p<0.01, ***p<0.001, ****p<0.0001.

We found that density of GS+ astrocytes was unchanged in zQ175 mice compared to WT mice, in both data from dm, vm and vl that was averaged together (whole striatum density) or when comparing within specific striatal regions (**Fig. 1C, D, E**). On the other hand, S100B+ astrocytes exhibited increased density in zQ175 mice compared to WT in the whole striatum (**Fig. 1F**), which was apparent in all three analyzed subregions (**Fig. 1G, H**), suggesting a homogenous increase in S100B+ astrocytes throughout the striatum in zQ175. This was consistent with previous reports indicating that neurodegeneration in HD occurs throughout the entire striatum (Bates et al., 2015; Morigaki & Goto, 2017). GFAP+ astrocytes also showed increased density in the whole striatum in zQ175 when compared with WT (**Fig. 1I**). However, when looking at GFAP+ astrocytes by different striatal areas in zQ175 mice, the density of GFAP+ astrocytes was significantly increased only in the dorsomedial striatum (**Fig. 1I, J, K**), a striatum region most affected in HD (Vonsattel et al., 1985), which accounted for the total increased density when looking at the whole striatum (**Fig. 1I**). These results showed a unique spatial distribution with GFAP+ astrogliosis only occurring in the dorsomedial striatum. Taken together, this data suggested a differential response of different astrocyte subtypes in both density and distribution throughout the striatum in zQ175 mice.

### The dorsomedial and dorsolateral striatum preferentially showed increased GFAP signal in zQ175 mice

While our cell density analyses showed a potential difference in the striatal distribution across different astrocyte subtypes, the obtained data was limited to the microscopy image area for any given striatal subregion which does not entirely represent the total area for each of the assigned striatum regions. To better estimate whether there were potential striatal regional differences in astrogliosis in HD, we utilized a macro zoom microscope to capture entire coronal slices. This type of imaging allowed for the visualization of fluorescence corresponding to each of the astrocyte markers in the entire striatum as well as other brain regions but did not allow for cell density analyses due to the limited resolution. Therefore, we decided to focus our analysis on the fluorescence intensity for GS, S100B, and GFAP in WT and zQ175 mice as proxy for astrocyte density and astrogliosis.

We used a systematic brain compartmentalization approach to define the area of different brain regions from where fluorescence intensity was measured and we sampled from the corpus callosum (CC), cortex (Ctx), and whole striatum (Str) (**Fig. 2A**). We also subdivided the striatum into four quadrants corresponding, relatively, to vm, vl, dm, and dl (dorsolateral) (**Fig. 2A**). We found that GS fluorescence intensity and S100B fluorescence intensity was unchanged between WT and zQ175 mice in all brain regions (**Fig. 2B-E**). The lack of changes in GS intensity between WT and zQ175 was consistent with the data obtained for GS+ density analyses (**Fig. 1B-D**). However, for S100B, the fluorescence intensity did not correspond to the changes observed in S100B density (**Fig. 1E-G**). A potential explanation would be the inability of the macro zoom microscope to capture fluorescence intensity changes derived from small nuclear areas, as would be seen with S100B. On the other hand, although GFAP fluorescence intensity was unchanged in the CC, it was significantly increased in the Ctx and Str in zQ175 mice compared to WT (**Fig. 2F, G**). Striatum degeneration is considered the major factor that influences HD symptomatology, but cortical degeneration is also prominent HD, especially in more advanced stages (ref). Within the striatum, we observed that GFAP fluorescence intensity was specifically increased in dl and dm striatum in zQ175 mice compared to WT, while no changes were observed in vm or vl striatum (**Fig. 2F, H**). Taken together, this data provided further evidence for regionally specific GFAP+ astrogliosis in HD.

**Fig. 2.**
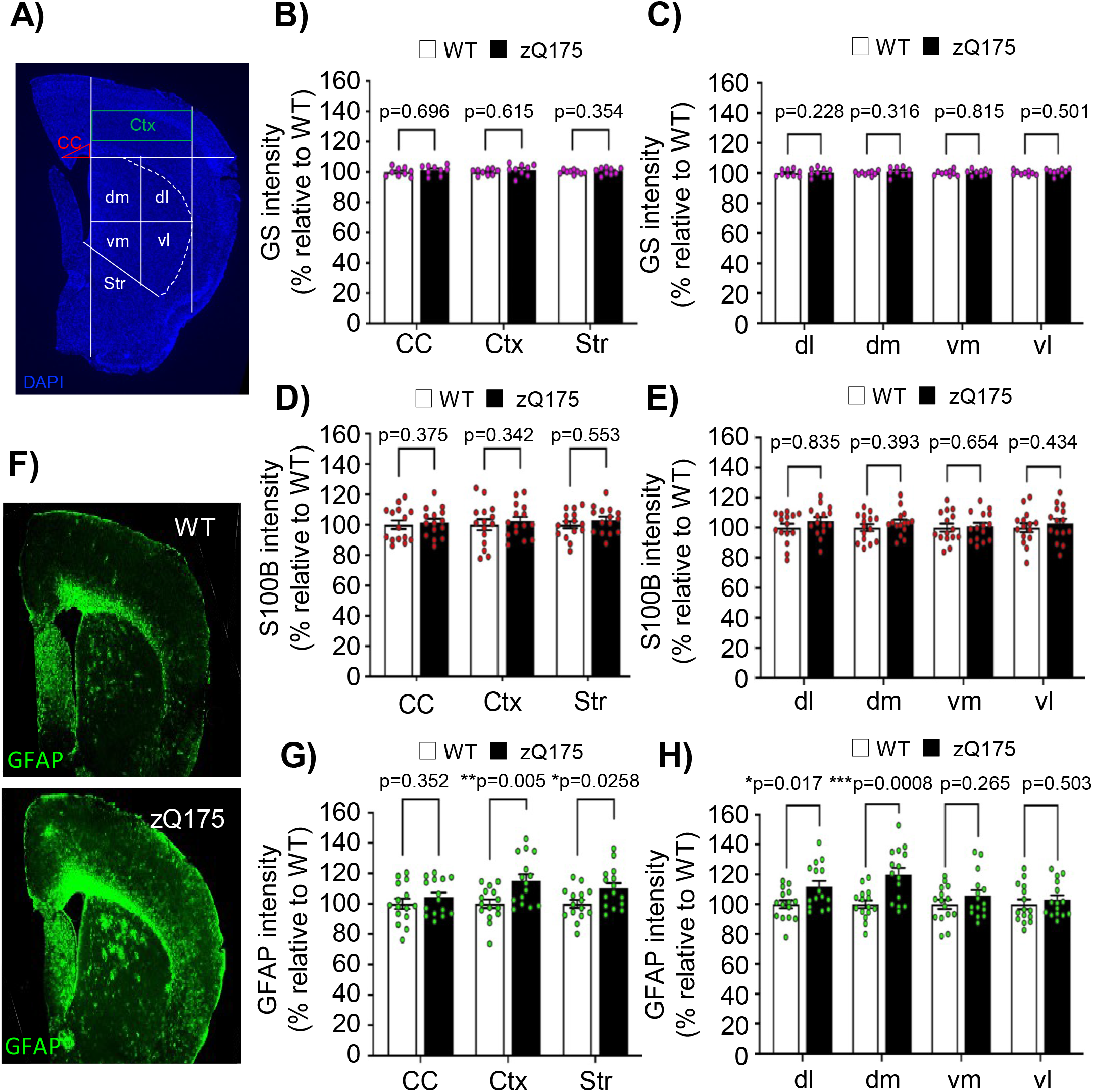
GFAP+ astrocyte fluorescence intensity is preferentially increased in the cortex and dorsal striatum of zQ175 mice. (**A**) Systematic brain compartmentalization approach used for fluorescence intensity analyses in the Cortex (Ctx), Corpus Callosum (CC), Striatum (Str), and striatal quadrants (dl, dm, vm, vl). DAPI stains nuclei. (**B, C**) GS fluorescence intensity in WT and zQ175 brains (n=9 slices from n=3 mice/genotype). (**D, E**) S100B fluorescence intensity in WT and zQ175 brains (n=15 slices from n=5 mice/genotype). (**F**) Representative image of GFAP fluorescence at 1.6X zoom. (**G, H**) GFAP fluorescence intensity in WT and zQ175 brains (n=15 slices from n=5 mice/genotype). Fluorescence intensity was calculated using FIJI from WT and zQ175 at 12 months. Data was relativized to 100% WT. Error bars denote mean ± SEM. Un-paired Student’s t test. *p<0.05, **p<0.01, ***p<0.001, ****p<0.0001.

### Clustering of striatal GFAP+ astrocytes is increased in zQ175

Previous analyses suggested that astrocyte somata are evenly distributed throughout any given brain region and that their processes overlap only minimally, indicating that each astrocyte covers an exclusive territory of neuropil and maintain a constant distant with their corresponding neighbors (Grosche et al, 2013). However, reactive astrocytes have previously been shown to cluster around aggregates and sites of injury presenting a non-uniform distribution (Buffo et al., 2008; Kamphuis et al., 2012). Clustered astrocytes are associated with focal inflammation and/or neurodegeneration. They can coordinate with microglia to remove dying neurons, with astrocytes specifically engulfing the diffuse apoptotic bodies derived from distal dendritic branches of dying neurons (Damisah et al., 2020). With the increased regional-specificity and heterogeneous distribution of GFAP+ astrocytes within the striatum of zQ175 mice, we studied whether GFAP+ astrocytes also presented a clustering pattern.

To determine astrocyte clustering, we calculated the nearest neighbor distance (NND), which measures the shortest distance from the center of one cell to the center of its closest neighboring cell (**Fig. 3A**). We calculated the NND for every astrocyte within the striatum of both WT and zQ175 mice and relativized the average NND to 100% in WT mice. We found that the NND for GFAP+ astrocytes in zQ175 mice was significantly lower (~30%) than in WT mice, and therefore were more clustered (**Fig. 3B, C, D**). To discard those changes in NND were due to increased density of astrocytes seen in zQ175 mice (**Fig. 1I-K**), we calculated the spacing index (NND^2^*density), which factors in density and therefore accounts for differences in cell density. Spacing index was significantly decreased in zQ175 mice compared to WT (**Fig. 3D**), further confirming that GFAP+ astrocytes were more clustered in zQ175 mice. Therefore, not only we observed increased density and specialized striatal compartmentalization of GFAP+ astrocytes but also increased clustering, suggesting the presence of specific foci within the striatum with enhanced neurodegeneration.

**Fig. 3.**
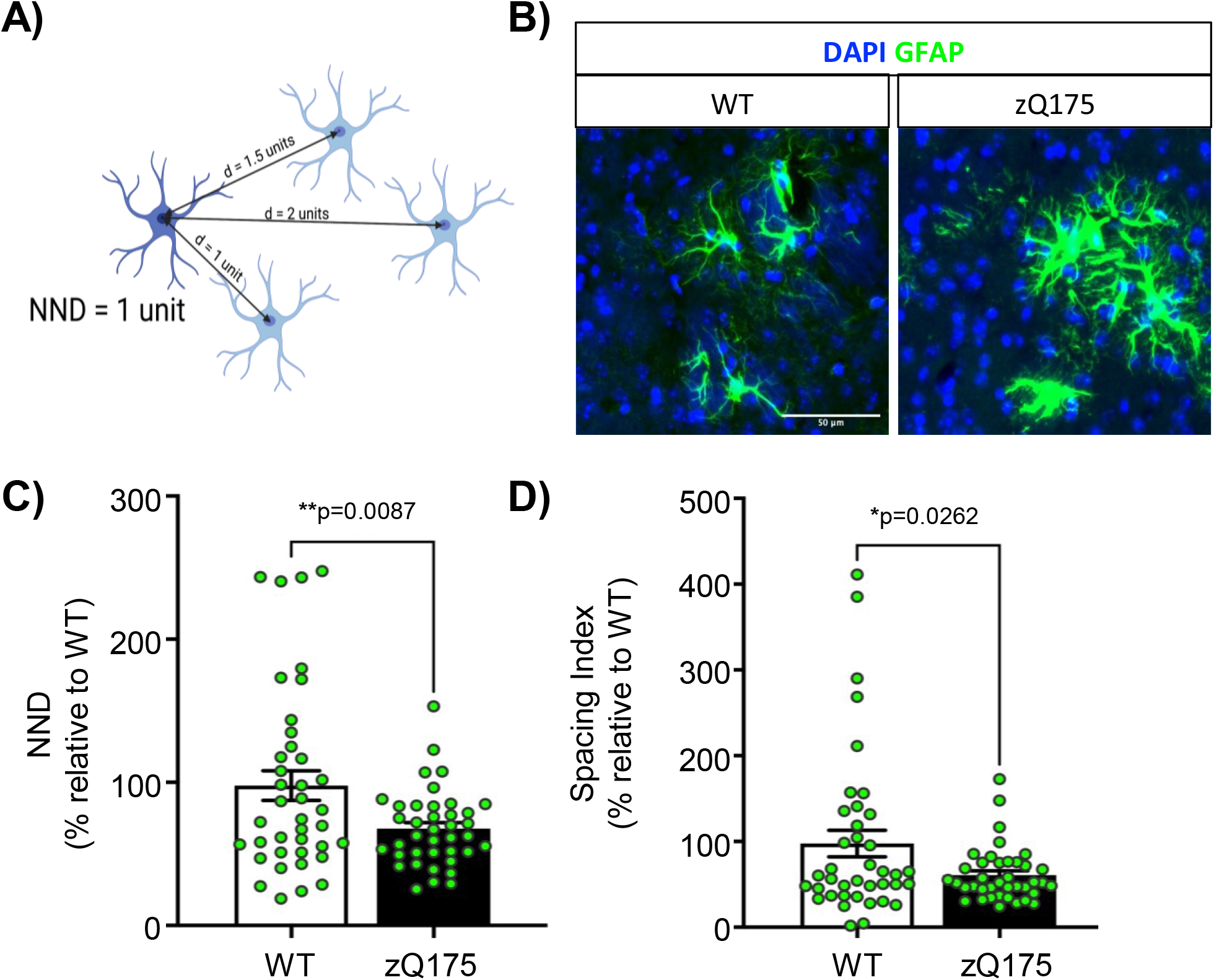
GFAP+ astrocytes in zQ175 mice present increased clustering. (**A**) Diagram of nearest neighbor distance (NND) analysis. Created on BioRender.com. (**B**) Representative immunostaining for GFAP (green) in the striatum of WT and zQ175 mice at 12 months. DAPI stains nuclei. Scale bar, 50 mm. (**C**) NND between GFAP+ astrocytes was calculated and relativized to WT (n=39 slices from n=13 mice/genotype). (**D**) Spacing index (NND^2^*density) between GFAP+ astrocytes were calculated and relativized to WT (n=39 slices from n=13 mice/genotype). Error bars denote mean ± SEM. Un-paired Student’s t test. *p<0.05, **p<0.01, ***p<0.001, ****p<0.0001.

## Discussion

Numerous studies have addressed the importance of astrocytes in HD (Al-Dalahmah et al., 2020; Behrens et al., 2002; Diaz-Castro et al., 2019; Faideau et al., 2010; Khakh et al., 2017; Lee et al., 2013; Lievens et al., 2001; Wood et al., 2019; Yu and Zarate et al., 2020). However, our understanding of the differential susceptibility of different astrocyte subtypes in HD was limited. We conducted a simultaneous comprehensive density and spatial analysis of multiple subtypes of astrocytes (GS, S100B, and GFAP) in symptomatic zQ175 mice. We demonstrated that different astrocytes subtypes respond differently under advanced disease conditions in terms of astrogliosis, spatial distribution, and clustering. Our data not only provided crucial information for understanding astrocyte pathology in HD but also highlighted the importance of proper astrocyte marker selection and accurate definition of striatum regions of interest when studying astrocyte biology in HD.

We have used three astrocyte markers in our study - GS, S100B, and GFAP. These markers were chosen based on a previous scRNA-seq study in which each of these markers was specific to one transcriptomic cluster of astrocytes in HD (Al-Dalahmah et al., 2020). A pertinent caveat of this analysis is the probabilistic overlap between subtypes. These subpopulations of astrocytes are not entirely distinct, but rather probably belong on a spectrum of expression, as they can exhibit partial colocalization (Du et al., 2021; Klein et al., 2020; Mack et al., 2018). While we acknowledge that partial overlapping between these three different markers may exist, our study focused on evaluating all astrocytes that were either GS+, GFAP+ or S100B+ and therefore changes in their overall density and/or distribution were not influenced by any partial colocalization. Future analyses will be necessary to investigate whether overlapping of one of more markers further establishes additional subtypes of astrocytes with distinct roles and susceptibility in HD.

GS (*Glul1*) is a pan-astrocyte marker in the striatum, as it is involved in the glutamate-glutamine shuttle and supplying neurons with glutamine for GABA and glutamate production (Hertz, 1979; Walls et al., 2015). Although glia pathology is consistent between striatal astrocytes from different mouse models of HD and those derived from patients with HD (Benraiss et al., 2021; Diaz-Castro et al., 2019), there is a lot of discrepancy about the directionality of GS levels. Studies in R6/1 and R6/2 HD mice showed that GS mRNA and immunoreactivity decreased respectively compared to WT mice, which was interpreted as a sign of excitotoxicity (Behrens et al., 2002; Khakh et al., 2017; Liévens et al., 2001). On the contrary, studies in zQ175 mice showed increased striatal density of GS+ astrocytes with disease progression (Yu and Zarate et al., 2020). However, although our study only compared WT and zQ175 at one symptomatic time point, we observed no changes in the density of GS+ astrocytes consistent with the absence of changes in GS protein levels reported in symptomatic BACHD (Lee et al., 2013). While we do not have an explanation for the discrepancies of GS levels and GS+ density across different mouse models, it is reasonable to hypothesize that the regulation of this gene marker can be influenced by factors other than mHTT.

Regarding S100B, its functions range between cell survival, protein phosphorylation, cytoskeletal dynamics, and intracellular Ca^2+^ homeostasis (Michetti et al., 2019, p. 100). S100B is increased in several neurodegenerative diseases, where it has been postulated to be induced by microglial cytokines and to cause both intracellular and extracellular effects that lead to neurotoxicity (Mrak & Griffin, 2001; Sathe et al., 2012; Serrano et al., 2017; Sheng et al., 1994). We found S100B+ astrocytes to be homogeneously distributed throughout the striatum with an increased density in zQ175 mice compared to WT. The homogeneity of S100B+ related astrogliosis is consistent with the widely accepted idea that striatal degeneration occurs throughout the striatum. Interestingly, we did not see this change reflected in our S100B intensity analysis. One potential explanation is the fact that despite having more cells expressing S100B in zQ175 mice the total protein levels throughout the striatum were not increased. Alternatively, the pixel resolution of the macrozoom microscope may not allow for the detection of the small, sparse areas of fluorescence we see with S100B. Nevertheless, the enhanced density of S100B+ astrocytes were consistent with increased striatal inflammation, as previously reported (Serrano et al., 2017).

GFAP marks intermediate filaments in astrocytes, and it is important for motility and structural stability of the cell (Eng et al., 2000; Hol & Pekny, 2015). Many studies point to GFAP+ astrocytes as a reactive subtype that is particularly enhanced in response to injury or inflammation (Buffo et al., 2008; Escartin et al., 2021; Hol & Pekny, 2015; Kamphuis et al., 2012). Recent studies support the functional enrichment of GFAP and S100B genes in HD, which are seen in different HD mouse models R6/2 and zQ175 and in human-derived cells (Benraiss et al., 2021). However, past HD studies are inconsistent in their use of GFAP as a marker for striatal astrogliosis due to this marker’s patchy pattern in the striatum. Our data demonstrated that GFAP ‘patchy’ immunostaining is indeed characteristic of GFAP+ striatal astrocyte pathology and that it is consistently enhanced in specific regions of the striatum. Additionally, we showed that GFAP+ astrocytes increased their clustering in zQ175 mice. We propose that GFAP+ astrocyte clustering may indicate specific areas that enhance astrocyte recruitment, such as areas of focal inflammation, synaptic degeneration, or MSN death. Further studies into the significance of these GFAP+ astrocyte clusters are needed. Nonetheless, our data demonstrate GFAP+ astrogliosis preferentially occurs in the most dorsal striatal regions and also in the cortex, a brain region that presents neurodegeneration at later stages of disease (Hedreen et al., 1991; Heinsen et al., 1994; Waldvogel et al., 2015).

In addition, we showed regionally specific astrocyte subtype changes within the striatum of zQ175 mice. Enhanced GFAP+ astrocyte density was observed in the dm and dl regions of the striatum. A potential explanation for such localized accumulation of GFAP+ astrocytes could be its association with neurodegeneration and/or behavioral alterations. MSN loss is widely reported in postmortem brains of patients with HD (Vonsattel et al., 1985), although no significant changes in the number of MSNs is observed in zQ175 or other mouse models of HD (Naver et al., 2003; Sun et al., 2002; Yu and Zarate et al., 2020). Therefore, accumulation of GFAP+ astrocytes within the dm and dl striatum may not be directly related to cell death. A recent study that characterized the excitatory input map of the striatum found that the dm and dl domains relate to goal-directed and habitual behaviors, respectively (Hunnicutt et al., 2016; Lipton et al., 2019). Interestingly, a reduction in goal-directed behaviors, which is associated with apathy, and altered habitual behaviors are common psychiatric symptoms in HD (Thompson et al., 2002). Therefore, it is possible that accumulation of GFAP+ astrocytes within the dm and dl striatum contribute to these behavioral alterations. Further studies to address the causality of GFAP+ clustering and astrogliosis in the dm and dl striatum onto behavioral deficits are warranted.

In summary, our study provided new insights on the differential alteration, both at the level of density and distribution, across different astrocytes subtypes in the brain of the zQ175 mouse model. Future studies in other animal models, in three dimensions, and in a longitudinal manner are needed for a full understanding of the dynamics of astrocyte heterogeneity and their differential susceptibility in HD.

## Methods

### Mouse lines

For this study we used a full-length knock-in mouse model of HD known as zQ175, which harbors a chimeric human/mouse exon 1 carrying an expansion of ~188 CAG repeats and the human poly-proline region (Menalled et al., 2012). WT (C57BL/6) animals were used as controls. Animals were analyzed at 12 months of age. Sample size was set to a minimum of three animals per genotype for every analysis. When possible, a balanced number of males and females were used. No randomization of animals was used in this study. All animal care and sacrifice procedures were approved by the University of Minnesota Institutional Animal Care and Use Committee (IACUC) in compliance with the National Institutes of Health guidelines for the care and use of laboratory animals under the approved animal protocol 2007-A38316.

### Immunohistochemistry

Mice were anesthetized with Avertin (250 mg/kg Tribromoethanol) and perfused intracardially with tris-buffered saline (TBS) (25mM Tris-base, 135mM Nacl, 3mM KCl, pH 7.6) supplemented with 7.5 mM heparin. Brains were dissected, fixed with 4% PFA in TBS at 4 °C for 4-5 days, cryoprotected with 30% sucrose in TBS for 4-5 days and embedded in a 2:1 mixture of 30% sucrose in TBS:OCT (Tissue-Tek), as previously described (Gomez-Pastor et al., 2017). Brains were cryo-sectioned at 16 µm-thick coronally, washed and permeabilized in TBS with 0.2% Triton X-100 (TBST). Coronal sections were acquired at roughly +1.5 to −0.3mm distance from Bregma. Sections were blocked in 5% normal goat serum (NGS) in TBST for 1 h at room temperature. Primary antibodies were incubated overnight at 4 °C in TBST containing 5% NGS. Secondary Alexa-fluorophore-conjugated antibodies (Invitrogen) were added (1:200 in TBST with 5% NGS) for 1h at room temperature. Slides were mounted in ProLong Gold Antifade with DAPI (Invitrogen), and fluorescent images acquired on an epi-fluorescent microscope (Echo Revolve). Primary antibodies used and dilutions are as follows: GFAP (Rabbit, Invitrogen PA1-10019, 1:500), GS (Mouse, BD Biosciences 610517, 1:200), and S100B (Rabbit, Abcam ab41548, 1:500).

### Imaging and Image Analysis

For cell number (GFAP, GS, S100B) the Cell counter plugin from ImageJ software was used and cells were counted manually. For intensity (GFAP, GS, S100B), the brain was systematically sectioned as shown in Figure 2. This was accomplished by creating two vertical lines going through the most dorsomedial portion of the lateral ventricle and the most lateral portion of the corpus callosum. A horizontal line was also drawn at the most dorsomedial point of the lateral ventricle. A fourth line was drawn from the most ventral tip of the lateral ventricle to the most ventral visible point of the corpus callosum. The four lines plus the corpus callosum were the boundaries of the striatum for measurement purposes. A triangle at the intersection of two perpendicular lines above the ventricle was used to sample corpus callosum intensity. The cortex was sampled from a rectangle between two vertical lines. The striatum was sectioned into four quadrants by calculating the maximum length and width and drawing horizontal and vertical lines at the midpoint. The fluorescence intensity density and area were measured for each brain region and striatal quadrant using ImageJ software. Intensity data was divided by the area of each individual region. For clustering (GFAP), the Nearest Neighbor Distance (NND) plugin on ImageJ was used. For all imaging experiments, 3 slices per animal were used.

### Experimental Design and Statistical Analyses

Data are expressed as Mean ± SEM, analyzed for statistical significance, and displayed by Prism 9 software (GraphPad, San Diego, CA). Normal distributions were compared with t-test (two-tailed) or ANOVA with appropriate post-hoc tests (Sidak’s, Dunn’s or Tukey’s) for multiple comparisons. Non-normal distributions were compared with the non-parametric Kruskal-Wallis test with appropriate post-hoc test, as indicated. The accepted level of significance was p ≤ 0.05, therefore samples indicated as n.s (no significant) did not reach p ≤ 0.05.

## Author contributions

R.G.P obtained funding for the study and designed the experiments. N.Z harvested mice used in the study. T.G.B. performed IF and imaging. T.G.B and M.T. analyzed the data. T.G.B. wrote the first draft of the manuscript and all authors edited subsequent versions. All authors read and approved the final version of the manuscript.

## Funding

This work was supported by R.G.P’s startup funds from University of Minnesota, the Biomedical Research Awards for Interdisciplinary New Science BRAINS (to R.G.P) and the National Institute of Health NINDS (R01 NS110694) (to R.G.P).

## Data Availability

All data generated in this study are presented in the current manuscript.

## Declaration of interest

The authors declare no competing interests.

## Notes

### Competing Interest Statement

The authors have declared no competing interest.

